# Sedation modulates fronto-temporal predictive coding circuits and the double surprise acceleration effect

**DOI:** 10.1101/2020.02.21.959171

**Authors:** Adrien Witon, Amirali Shirazibehehsti, Jennifer Cooke, Alberto Aviles, Ram Adapa, David K. Menon, Srivas Chennu, Tristan Bekinschtein, Jose David Lopez, Vladimir Litvak, Ling Li, Karl Friston, Howard Bowman

**Affiliations:** School of Computing, University of Kent, Kent, United Kingdom, CT2 7NF; East Kent Hospitals University NHS Foundation Trust, Kent & Canterbury Hospital, CT1 3NG; Institute of Psychiatry, Psychology & Neuroscience, King’s College London, London, United Kingdom, SE5 8AF; Division of Anaesthesia, Box 97, Cambridge Biomedical Campus, University of Cambridge, CB2 0QQ, UK; Department of Psychology, University of Cambridge, Cambridge, CB2 3EB, UK; Electronic Engineering program, Universidad de Antioquia, Ciudad Universitaria, Medellín, Antioquia, Colombia; Wellcome Centre for Neuroimaging, University College London, London, UK; School of Psychology, University of Birmingham, Birmingham, B15 2TT, UK; Center for Neuroprosthetics, EPFL Valais, Sion, Switzerland

## Abstract

Two important theories in cognitive neuroscience are predictive coding and the global workspace theory. A key research task is to understand how these two theories relate to one another, and particularly, how the brain transitions from a predictive early state to the eventual engagement of a brain-scale state (the global workspace). To address this question, we present a source-localisation of EEG responses evoked by the local-global task – an experimental paradigm that engages a predictive hierarchy, which encompasses the global workspace. The results of our source reconstruction suggest three-phases of processing. The first phase involves the sensory (here auditory) regions of the superior temporal lobe and predicts sensory regularities over a short timeframe (as per the local effect). The third phase is brain-scale, involving inferior frontal, as well as inferior and superior parietal regions; consistent with a global neuronal workspace (as per the global effect). Crucially, our analysis suggests that there is an intermediate (second) phase, involving modulatory interactions between inferior frontal and superior temporal regions. Furthermore, sedation with propofol reduces modulatory interactions in the second phase. This selective effect is consistent with a predictive coding explanation of sedation, with propofol acting on descending predictions of the precision of prediction errors; thereby constraining access to the global neuronal workspace.

## Introduction

Two important theories in current cognitive neuroscience are predictive coding [Rao1999, Friston2010] and global neuronal workspace theory [Dehaene2011]. The former emphasizes forward and backward exchanges along sensory processing and higher level pathways, with forward connections carrying prediction errors, and backwards connections conveying predictions. In contrast, the latter emphasizes a distinct mode of processing – the global workspace – which has the character of a sustained brain-scale state, into which there is a sharp transition – described as *ignition* [Dehaene2006].

Indeed, King et al. argued for the existence of two distinct modes of processing; the first restricted to sensory areas and the latter the global neuronal workspace. In addition, they have suggested that these two modes are experimentally engaged by the local-global task [King2014]. This is an auditory deviance task in which tones can be unexpected at two different levels of regularity. The first level, which generates the (so called) local effect, reflects regularity at a short temporal frame of reference; i.e. repeated *tones*. In contrast, the second level, which generates the (so called) global effect, reflects regularity at a longer temporal frame; i.e. repeated *sequences* of tones.

In this way, the local-global task engages a predictive coding hierarchy, with multiple levels at which prediction errors could arise, in much the same vein as proposed by predictive coding [Friston2006]. Global workspace theory would, though, additionally propose the existence of a spatially broad and temporally extended “brain-scale” state at the top (or centre) of this hierarchy, which would be associated with the global effect. Key to reconciling predictive coding and global workspace theory is to understand how the transition from a predictive early stage (disclosed by the local effect) to engagement of a brain-scale state (disclosed by the global effect) is mediated. Understanding this transition – or ignition – is the objective of the work presented here. In other words, how do lower levels of a processing hierarchy come to engage higher levels – and what are the underlying neurophysiological mechanisms? In this respect, our work contrasts interestingly with recent research with related objectives, e.g. [Chao et al, 2018].

Predictive Coding (PC) postulates relatively fast coordinated exchanges up and down a multi-layered hierarchy, while the Global Neuronal Workspace (GW) argues for longer term, “meta-stable” states; clearly, how one transitions between these two is of fundamental interest. Additionally, in terms of similarities, both predictive coding and the global workspace assume an underlying hierarchy of neuronal message passing or what, from a Bayesian perspective, could be considered belief propagation. Furthermore, both associate higher (or deeper) levels with perceptual synthesis at longer timeframes.

Under predictive coding, the influence of ascending prediction errors depends upon their precision (i.e., reliability or inverse variance). This has often been cast in terms of attentional selection, where ascending prediction errors that are afforded greater precision are selected to have a greater influence on belief updating at higher hierarchical levels (Auksztulewicz and Friston 2015, Kanai, Komura et al. 2015, Parr and Friston 2017). On this view, ignition – or a transition to global processing throughout the depth of cortical hierarchies – rests upon the top-down control of the precision of bottom-up signals (i.e., prediction errors). This gracefully relates attentional processing to conscious content that gains access to the global workspace, while referring to a measurable aspect of neuronal processing; namely, the modulation of excitability of synaptic gain (which precision controls) of neuronal populations reporting prediction errors.

One clear hypothesis that follows from predictive coding, and has informed our study, is that a transition from local to global processing would be accompanied by descending modulation from higher cortical levels to lower cortical levels. To inform this hypothesis empirically, we use the local global paradigm – in conjunction with source localisation (to establish the hierarchical level of neuronal activity) – and characterise responses evoked by (local and global) violations as a function of peristimulus time. To test the hypothesis that evoked responses reflect modulatory interactions between high and low cortical levels, we crossed the local global paradigm with an inhibitor of neuromodulation; namely, propofol. Propofol has been previously shown to modulate extrinsic hierarchical connections, i.e. involving across-brain sources (Boly, Moran et al. 2012, Gómez, Phillips et al. 2013), and indeed, to do this within the context of the local-global task [Uhrig et al, 2016; Nourski et al, 2018]. Furthermore, the use of an anaesthetic offers the opportunity for construct validation; in the sense that it reduces levels of conscious processing, of the sort associated with the global workspace (Uhrig, Janssen et al. 2016),

Our previous work with the local-global task (focusing on responses in sensor space), suggested the transition from local to global processing is not temporally or functionally as sharp as one might have thought [Shirazi2018]. Indeed, our findings could be interpreted as evidence for a transitional phase between the local and global responses. This evidence rests on an interaction between the local and global conditions [Shirazi2018]. In other words, early responses to global violations depend on the presence of a local violation. Alternatively, the late responses to local violations depend upon a global violation. This interaction appears to be driven by changes in responsiveness during the positive rebound to the N1 and mismatch negativity (MMN); that is, during the P3a window (∼200ms to ∼400ms post stimulus onset). The key aspect of this change in responsiveness is a coincidence of surprise; i.e., a combination of local and global deviants induces an acceleration of the P3, an effect we have called the *double-surprise acceleration*^**1**^ *effect* [Shirazi2018]. Furthermore, manipulation of awareness (through sedation) was found to modulate this interaction.

In this paper, we extend these findings. As just discussed, we have established a transition between local and global processing phases, temporally and functionally. Here, we characterise this transition neurophysiologically – in terms of hierarchical processing in the brain. That is, we ask the following question: can we – in respect of brain areas – observe a transition from a localised (low-level) prediction error to a (high-level) global ignition? In particular, can we neurophysiologically identify the phase transition, which we propose is an additional transitional stage? A further key question is the role of awareness in this hierarchical processing. In particular, as Dehaene et al. would argue; is it only the late, global processing that engenders states of awareness [Dehaene2011] (see also, Uhrig et al, 2016)?

To answer these questions, we report a Multiple Sparse Priors [Friston2008] source localisation of the local-global task. This enabled us to characterise the neurophysiological trajectory of neuronal responses as they propagate from rapidly changing early responses restricted to sensory areas (for us in auditory cortices) to slowly changing late (c.f., meta-stable) responses (involving temporal, frontal and parietal areas). Our key finding was that the transition between these two phases – sensory-bound to global – involves a transient engagement of the superior temporal–inferior frontal network. Furthermore, the interaction between local and global violations on responses in this network is attenuated by propofol sedation.

## Methods

### Participants

Originally, 22 neurologically healthy adults were included in the study, but two recordings were lost due to technical issues, leaving 20 participants (9 male; 11 female) (mean age = 30.85; SD = 10.98).

### Experimental Design

The local-global auditory oddball task, devised by Bekinschtein [Bekinschtein2009], was used to characterise differences between local and global effects after healthy sedation and subsequent recovery. As shown in Figure 1, local regularity was established using sequences of five tones, or quintuples, where the last tone may or may not vary from the preceding four tones (local deviant versus local standard respectively). Global regularity was established as the most frequently presented quintuple type within a block, either local standard (all five tones the same) or local deviant (different last tone). Thus, violations in global regularity were expressed by the presentation of a quintuple that differed from the frequently presented type in any block. To ensure global regularity was established, a habituation period of 20 to 30 quintuples was presented at the beginning of the block. After the habituation phase, the ratio between the standard and deviant quintuples was set to 80/20. This created four conditions: (1) local standard / global standard (LSGS), (2) local deviant / global standard (LDGS), (3) local standard / global deviant (LSGD) and (4) local deviant / global deviant (LDGD) (see b, c, d and a in Figure 1). Quintuples comprised 5 tones of 50ms duration each, presented via headphones, with an intensity of 70dB and an SOA of 150ms. All tones were synthesised with 7ms rise and 7ms fall times. Participants were asked to count the number of global deviants they heard during both sedation and recovery phases of the study as an incidental task to reduce fluctuations in attentional set.

**Figure 1.**
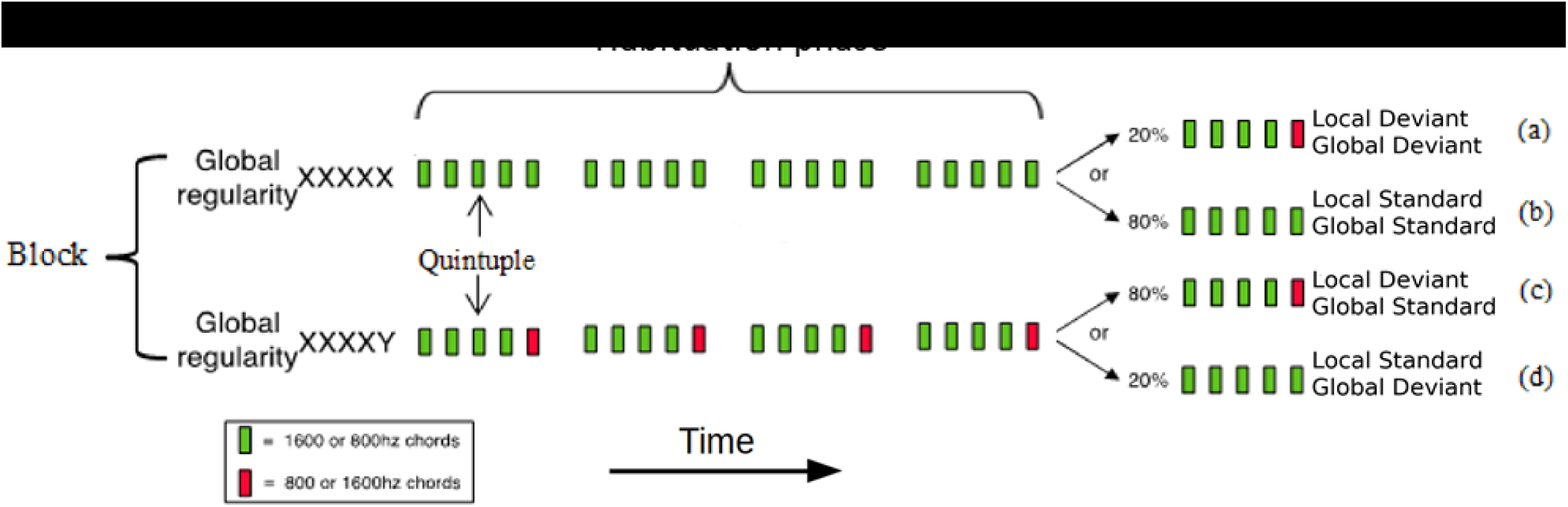
local-global auditory task design from [Bekinschtein2009].

### Sedation

During surgery or procedures for diagnosing medical conditions, it is common to take the patient to a sedative plane (as opposed to general anesthesia); also for pain control it is common to use propofol to relax the patient or take him/her/they to the point of sleep. In sedation research studies, the state is defined either by the target concentration in blood (light, medium or moderate sedation; light, medium or deep anesthesia) and /or a clinical responsiveness scale like the Ramsay^2^. However, in the study analysed in this paper, the sedation states are defined in an even more detailed manner, by exact concentration of propofol in blood in each state and by RTs and response misses in a behavioral task, as described for this experiment in [Chennu2016b].

In the experiment we are analysing, we knew that the target concentrations induced light to moderate sedation, where participants range from mild changes in relaxation to mostly unresponsive, but easily “arousable”, since these dose-responses had been defined already, as in [Stamatakis2010, Adapa2014, Barttfeld2015, David2007].

Specifically, we knew the behavioral and drug in blood pattern, enabling us to place participants around “the verge of unconsciousness”. The specifics of how we manage to maintain participants in this state, with such small doses are elaborated in [Absalom2009].

The local-global task was presented on two occasions; once during either mild (half of participants) or moderate (the other half) sedation and once 20 minutes later, when participants were considered to be in recovery (i.e. no longer sedated). Sedation in this study induced a heavily relaxed but behaviourally responsive state. All participants were tested both under sedation and subsequently in recovery, creating a repeated measures design. Each experimental run began with an awake baseline period lasting 25-30 minutes followed by a target-controlled infusion of propofol [Marsh1991], administered via a computerized syringe driver (Alaris Asena PK, Carefusion, Berkshire, UK). Three blood plasma levels were taken– μg/ml (mild sedation), 1.2 μg/ml (moderate sedation), and after recovery from sedation. A period of 10 minutes was allowed for equilibration of calculated and actual plasma propofol concentrations before cognitive tests commenced. Following cessation of infusion, plasma propofol concentration exponentially declined toward zero and approached zero in 15 minutes leading up to behavioural recovery. In light of this, the recovery condition started 20 minutes after cessation of sedation.

All procedures were conducted in accordance with the Declaration of Helsinki. The participants provided written informed consent and were healthy controls. Ethical approval for testing healthy controls was obtained from the Cambridgeshire 2 Regional Ethics Committee.

### EEG recording

During pre-processing, two patients were excluded due to artefacts, therefore 18 participants were taken forward for analysis. Participants were asked to close their eyes during data collection to avoid eye artefacts in the data. EEG data were collected on two occasions: during sedation and then recovery. A Net Amps 300 amplifier (Electrical Geodesic Inc., Oregon, USA) with a high-density cap of 129 channels was used for data acquisition; and preprocessed data were obtained using custom Matlab scripts based on EEGLab. EEG timeseries were recorded in microvolts (μV), with a sampling frequency of 250Hz, and referenced to vertex (channel Cz). After recording, the data were segmented from −200ms before the first tone in a quintuple until 1296ms after that tone. Bad channels (those that crossed a 200uVs threshold) were interpolated. The remaining trials were re-referenced to their average and band-pass filtered from 0.5 to 20Hz – the standard filter settings for this paradigm [Bekinschtein2009]. Each dataset was then converted to SPM12 format (http://www.fil.ion.ucl.ac.uk/spm/) for subsequent analysis. Channels near the neck and eyes were discarded after conversion; since these are often sources of muscle and eye-movement artefacts (36 out of the 129 channels).

### EEG source reconstruction

All analyses focussed on the transient evoked by the fifth tone in a quintuple. Accordingly, trials were corrected to a baseline 200ms before the onset of the fifth tone, which occurred between 400ms and 600ms from trial (i.e. 1st tone) onset. The time segment used for analysis was −200ms (from fifth tone) to the end of the trial at 696ms from fifth tone onset. A group inversion was performed to minimize the variance in source localisation (over participants) using Multiple Sparse Priors (MSP) – the Bayesian inversion scheme within SPM [Litvak2008, Friston2008]. After inversion, three windows of interest were selected – as explained in the next section – to study the sources generating responses to the fifth tone. For each window, images of evoked power (i.e. squared deflection from zero), in source space, were computed for each condition and participant. The General Linear Model (GLM) was used to test for significant effects of the four conditions at each source. The ensuing statistical parametric maps (SPMs) in source space were corrected for multiple comparisons using Random Field Theory in the usual way.

In brief, the source reconstruction proceeds in three steps:

- A co-registration step over participants, which maps sensor-coordinates to source space (dipole) coordinates.
- A forward model for the transformation matrix (Lead-field) from dipole activity to scalp activity.
- The inverse transformation, which optimizes the solution at the source level to best explain scalp data (in terms of sparse sources).

Co-registration between the scalp level and the source level in MRI space was applied. Coordinates at the scalp level were based on a standard GSN-Hydrocel template with 128 channels. Three fiducials were used to map the coordinates from sensor space to source space: nasion, left peri-auricular point and right peri-auricular point. A template head model based on the MNI brain was applied with a cortical template mesh of 8196 dipoles, which contains the coordinates of the dipole sources.

As previously stated, we discarded 36 electrodes on the face, neck and cheek from the montage, since they were noisy and dominated by muscle artefacts. This left us with incomplete coverage, making the localisation problem more difficult. As a result, we constrained the cortical mesh by selecting cortical areas that are, a priori, most likely to generate evoked responses; i.e., only regions of the temporal, frontal and parietal lobes were included for source reconstruction. Figs. 9 and 10 and supporting text in the supplementary material provide a detailed justification for this choice. We did not include deep sources or sources in the occipital and motor cortices, as there is no prior evidence suggesting these regions are related to the effects of interest in the present study.

The forward model was computed using the BEM (Boundary Element Method) [Phillips2007] as standard in SPM12 for EEG-based source reconstruction, with a three layer head model, i.e. skin, skull, and brain. At the source level, a mesh based on an MRI template is used to simulate the *1484* dipolar amplitudes in the brain.

We selected a time-frequency window for the source reconstruction. The frequency band used was the same as used in pre-processing with a range of [0.5 20] Hz. The window used for the source reconstruction is from 400ms after the first stimulus onset in a quintuple, to the end of the analysed segment. This included the baseline and the evoked response associated with the fifth tone up to the end of the quintuple.

### Window placement for image extraction

After source inversion, two analyses were performed in order to characterise the spatial and temporal responses. The first used statistical parametric mapping to test for differences in evoked responses in all sources. The second analysis focused on responses in Regions Of Interest (ROI) based on regionally specific time-series at the source level. The ROIs were defined based upon effects identified by the statistical parametric mapping, as explained below.

Statistical parametric mapping summarizes the activity on the mesh, across the time window chosen. The evoked power was averaged within time windows for the frequency range [0.5 20] Hz. A spatial filter (FWHM=1mm) was used to smooth the dipole activity in three dimensional source space.

Evoked power under each condition was calculated as the root mean squared response (in source space) over the window. Statistical inference in this context, then, required the placement of time windows. Tailoring such windows post-hoc to the EEG data, risks biased sampling and could inflate false positive rates (e.g. [Brooks2016]). Consequently, prior precedents for these placements are used. These were taken from [Bekinschtein2009], which introduced the local-global experimental paradigm (see section “Window Placement in the supplementary material for further justification of these window settings). Specifically, we looked at the following three effects:

- Early window [100,150] ms: to quantify the local effect, which would be expected to correspond to the mismatch negativity in electrode space;
- Middle window [250,350] ms: to assess the interaction between the local and global effects, which is most likely to occur when both local and global effects are present;
- Late window [400 600] ms: to quantify the global effect, which would be expected to correspond to a P3b in electrode space.

## Statistical analysis

### Hierarchical spatial responses

For each participant (18) and each condition (8), three images of evoked power (one for each time window) were created for General Linear Model (GLM) statistical analysis. The experimental design can be summarized as a 2×2×2 within-subjects design, with 3 factors: sedation, local, and global. Each factor comprises 2 levels: sedation and recovery (for sedation); local standard (LS) and local deviant (LD) (for local); and global standard (GS) and global deviant (GD) (for global). The ensuing statistical analysis can be summarized as follows.

The first issue was to understand the relationship between local and global manipulations. To do so, we first looked at the local effect and the global effect individually and then the local x global interaction. Since the local and global effects have been extensively explored and are well documented in the literature; our analyses of these effects serve as sanity checks of our source localisation. That is, if the MSP algorithm localises these effects to the expected brain areas, we can have confidence that the reconstruction scheme can localise the effects for which there are fewer precedents.

The second issue was to understand the effect of sedation. We therefore assessed the main effect of sedation, the sedation x local interaction, the sedation x global interaction and the three way (sedation x local x global) interaction.

We conducted a flexible ANOVA analysis with pooled variance (see Supplementary Material section “Pooled Variance”), which employs a two-step threshold to control for multiple comparisons [Friston2007]. The first level cluster-forming threshold used an uncorrected alpha level of 0.001 to define clusters of voxels. Secondly, Random Field Theory was used to determine the likelihood that a cluster of voxels of a particular size will arise under the null hypothesis. The (cluster level) threshold was a Family Wise Error corrected alpha level of 0.05. This choice of thresholds, provides robust protection against false positive rates [Flandin,2016].

### Temporal dimension

Source analysis reveals which cortical sources exhibit significant effects of interest, within the different time windows. For illustrative (but not statistical) purposes, we investigated how activity changes through time at the source level, as follows. A source in each brain region – selected to plot the source time-series – was taken from the peak of the significant cluster, during the middle window [250, 350] ms. Specifically, we selected the temporal lobe sources located at the peak of the cluster for the local effect, the frontal lobe sources located at the peak of the cluster in the local x global interaction, and the parietal sources located at the peak of the cluster for the global effect.

The sources were then identified within an ROI of 5 mm and the corresponding time-series were exported for each subject and each condition. Since statistical results are not being reported on these time-series, issues of double dipping do not arise.

Only the left hemisphere time-series are presented for illustration (the time course for the right hemisphere was very similar). The regional timeseries data correspond to the group average of the time-series for each window; namely, the early window [50 100] ms, the middle window [250 350] ms and the late window [400 600] ms. The regional times-series were then preprocessed to provide a quantitative characterisation of effect sizes. The regional time-series were smoothed with an adapting hamming window: the points up to the end of the early window were smoothed with a hamming window width of 50 ms. The length of the hamming window then increased linearly from 50 to 100 ms up to the beginning of the middle window. Then, the window size remained the same until the end of the middle window. To deal with the second transition period, the size of the hamming window increased linearly from 100ms to 200 ms until the beginning of the late window. From the beginning of the late window onward, the hamming window length was fixed at 200 ms. (This adapting hamming window ensures that the effect size at the centre of each window is the value that is entered into the SPM inference). Finally, the root mean square was taken over the source time-series within each region of interest. To deal with edge effects, the data were mirrored in time at the beginning and the end of the epoch.

## Results

In this section, the results of the local effect, the global effect, the local-by-global interaction and the three way interaction are presented. Note that the condition specific time-series presented in this section are positive at all points; i.e., they report evoked power (i.e., root mean square responses).

### Local effect

The local effect (green line) is significant in the temporal sources, during both the early and middle windows, as shown in Fig. 2A) and B). The time course for the temporal area is shown in Fig. 2C) where the MMN appears clearly with a peak in the early window (dashed blue arrow). The local effect is again significant in the temporal region during the middle window (solid blue arrow), whereas it is not significant in the late window. Fig. 2B) shows that frontal clusters are also significant in the middle window. This is shown by the time course of the frontal cluster in Fig. 2D), with an effect peak during the middle window. Table 1 summarizes the statistical results for each cluster in the early and middle windows. For each cluster, the peak location is described in the second column and the F-statistic for the peak of the cluster in the third column. The family-wise error (FWE) corrected p value is presented (4th columns) with the size of the cluster (5th columns). This shows a strong effect (P_FWE_ < 0.001) for all the clusters shown in the table.

**Table 1.** Statistics for each cluster for the early and middle windows. Each cluster, named in the first column, is characterized by its peak location in MNI coordinates as shown in the second column, the F-value for that peak (third column), the p-value (fourth column) and the cluster size (last column). The p-value highlights the significant cluster after family-wise error correction, set to an alpha of 0.05.

| Clusters | Peak location<br>(x,y,z) | F[1,119] | P<sub>(FWE)</sub> | K cluster size |
| --- | --- | --- | --- | --- |
| Early window |  |  |  |  |
| Left temporal | (-58,-24,4) | 15.24 | <10<sup>-3</sup> | 383 |
| Right temporal | (52,-24,4) | 15.14 | 10<sup>-3</sup> | 312 |
| Middle window |  |  |  |  |
| Left temporal | (-58, -24, 4) | 21.02 | <10<sup>-3</sup> | 491 |
| Right temporal | (42, -20, 8) | 18.9 | <10<sup>-3</sup> | 365 |
| Left frontal | (-46, 20, -10) | 22.65 | <10<sup>-3</sup> | 591 |
| Right frontal | (-44, 22, -12) | 22.59 | <10<sup>-3</sup> | 584 |

**Figure 2.**
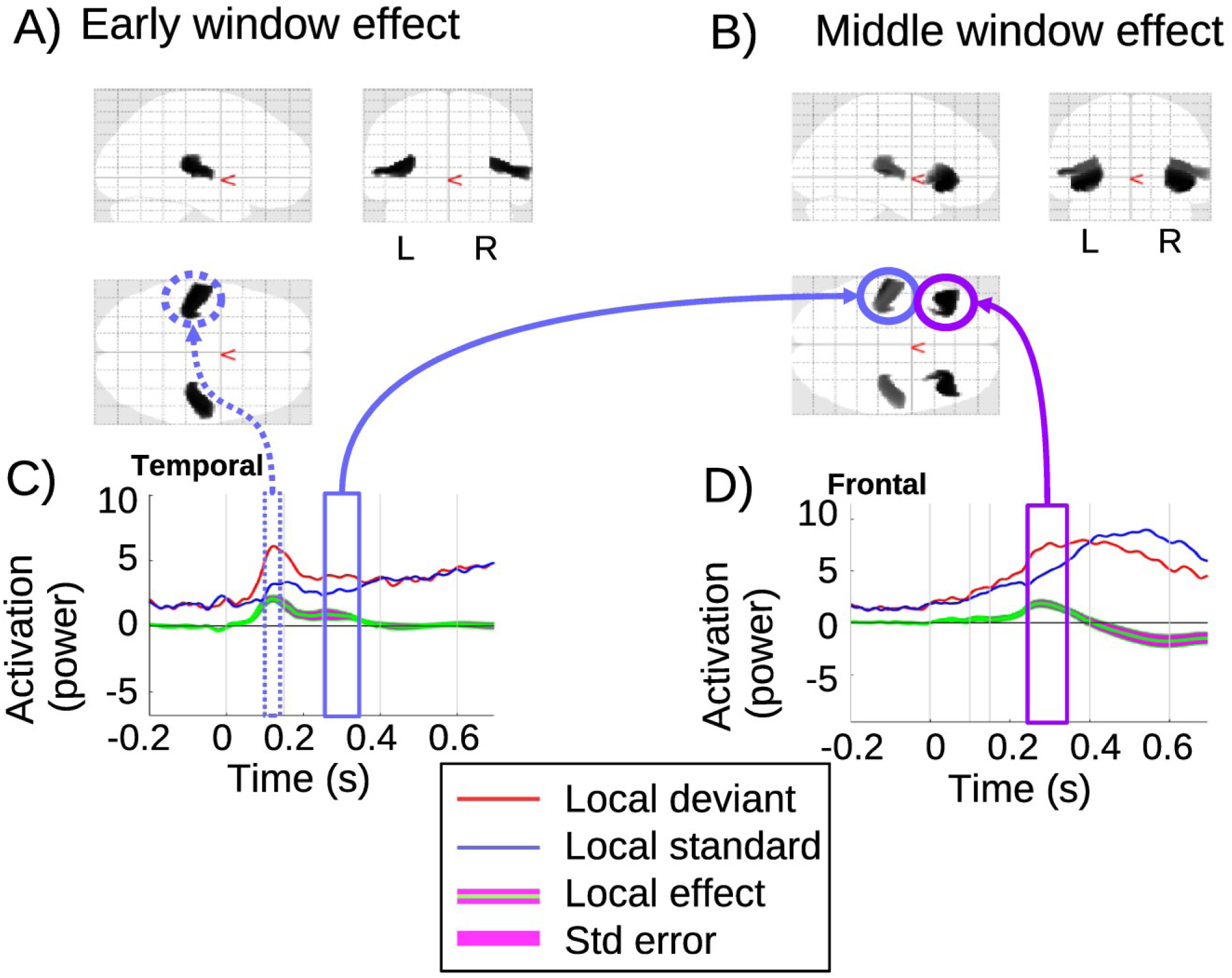
Local effect: A,B) present the SPM results with the significant clusters in a 3D glass brain image for the early (A) and middle (B) windows. C,D) The source time-series are plotted for the clusters in the temporal (C) and frontal (D) lobes. Zero is the onset of the (critical) fifth tone. The times-series are summarized across subjects and shown in red and blue for local deviant and local standard respectively. The local effect between the two conditions is plotted in green, and the standard error in magenta.

### Global effect

The global effect is presented in Fig 3. In the early window, the global effect is significant in both left and right frontal sources, as shown in Fig 3C). This early global effect can be related to the contingent negative variation (CNV) [Chennu2013], which is usually observed at frontal electrodes. Accordingly, the time course for the frontal area in Fig 3F) shows that the global deviant is greater than the global standard before and during the early window. During the baseline, a small global effect (before the onset of the fifth tone) is also consistent with this CNV effect, as an anticipation of the global deviant quintuple. The nature and implications of this CNV effect are discussed in Section S6 in the Supplementary Material of [Shirazi2018], where it also appears on the scalp.

**Figure 3.**
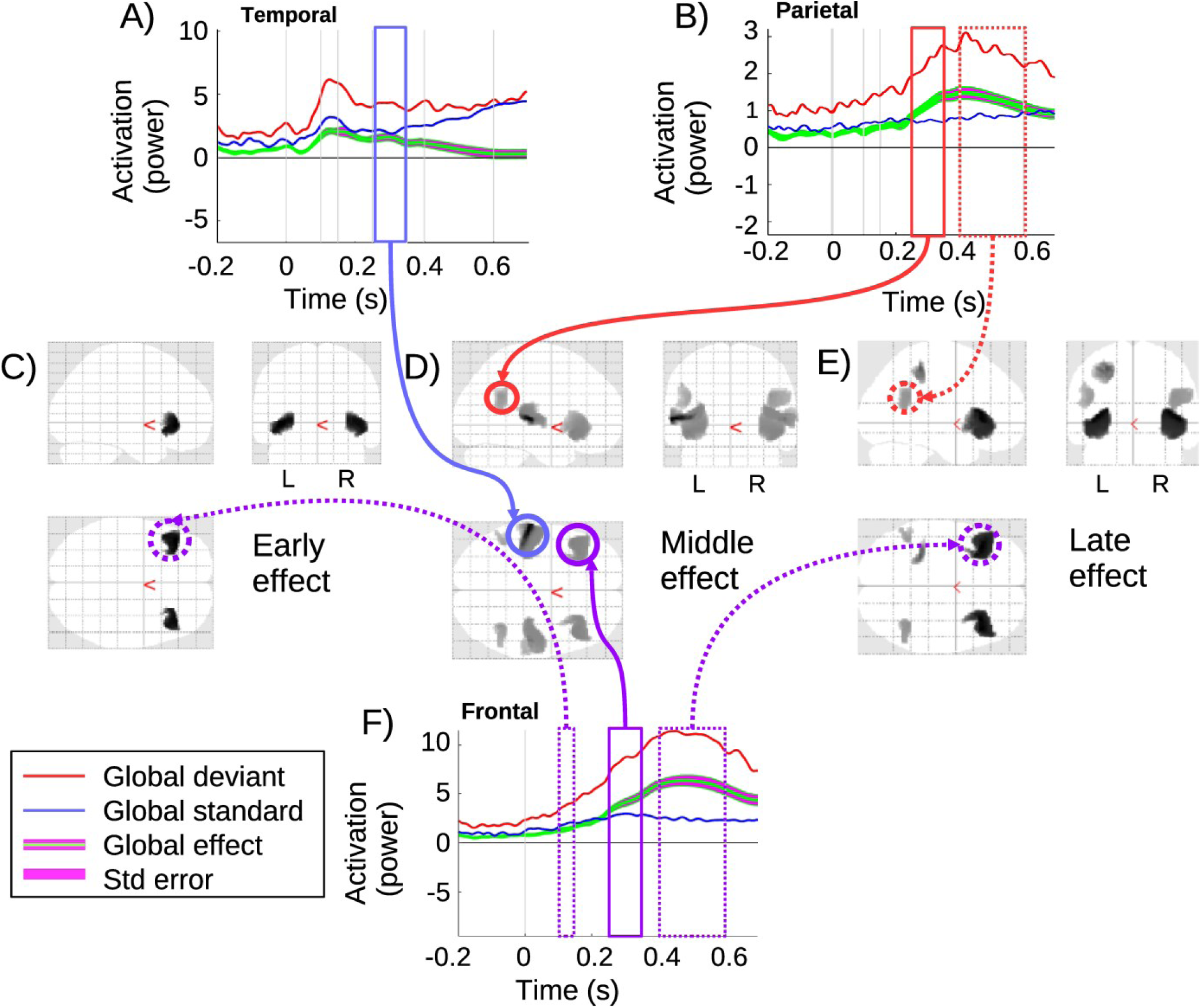
Global effect: A) source time-series corresponding to the temporal lobe cluster, with significant effect in the middle window; B) source time-series corresponding to the parietal cluster, with significant effect in middle and late windows. C,D,E) 3D glass brain images with significant clusters in early, middle and late windows. F) Source time-series at the frontal cluster, with significant clusters in all three windows. Zero is the onset of the (critical) fifth tone.

During the middle window, as shown in Fig 3D), the global effect is significant in the temporal, parietal and frontal regions with the time course represented respectively in Fig 3A), Fig 3B) and Fig 3F). Consistent with Wacongne et al 2011 and Chennu et al 2013, this window involves a broad network of brain activity indicated by the significant clusters. Fig 3A) shows the time course in the temporal area. The global deviant in this area diverges from global standard, but is significant only during the middle window, before disappearing in the late window.

In the late window, Fig 3E), the global effect is significant in a network comprising both frontal and parietal regions. The time course for the parietal cluster is plotted in Fig 3B), which shows the global effect is significant during the middle and late windows with a peak at the beginning of the late window. Additionally, specifically on the left side, a second more dorsal parietal cluster appears which was not present in the middle window. Finally, Fig 3F) shows the frontal time course of the global effect, which is significant in all three windows, with a peak in the middle of the late window. This frontal area is the most activated for the global effect, starting from the CNV, until achieving the strongest effect during the late window.

The statistical results from SPM for each significant cluster are shown on Table 2 below. The clusters in the early window have p-values (FWE-corrected) of approximately 0.01, while the strongest effects appear during the middle window, with p-values (FWE-corrected) below 0.001 for all clusters.

**Table 2.**
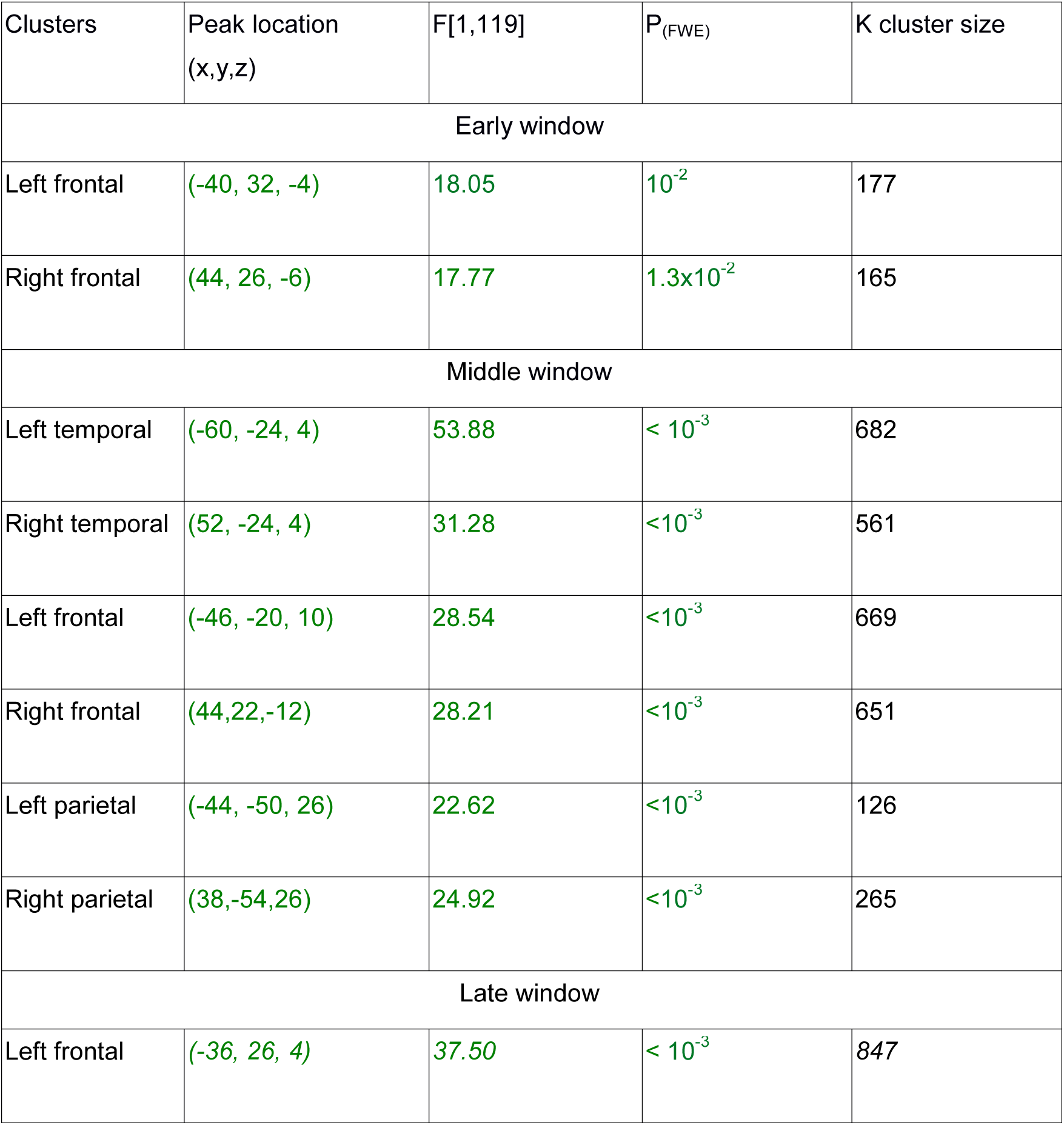
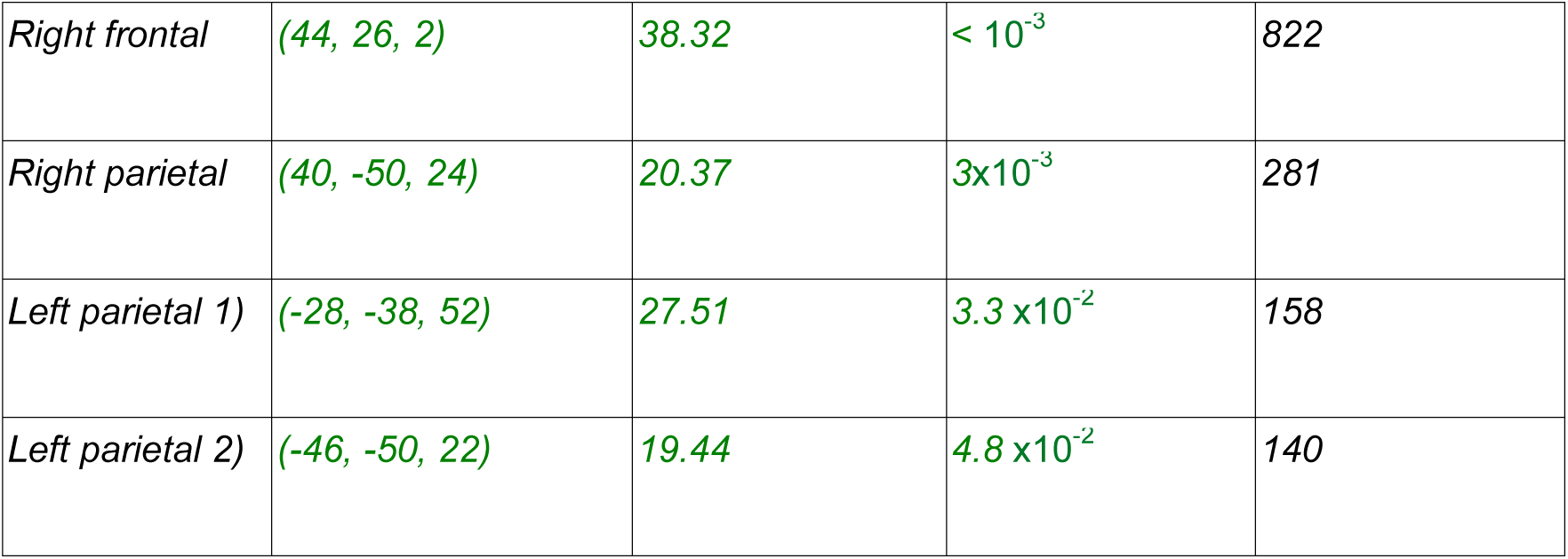
Statistics for each cluster for the early, middle and late windows. Each cluster, named in the first column, is characterized by its peak location in MNI coordinates, as shown in the second column, the F-value of the peak (third column), the p-value (fourth column) and the cluster size (last column). The p-value highlights the significant cluster after family-wise error correction, set to an alpha of 0.05.

### Local-by-global interaction

The local-by-global interaction is significant only in the middle window, as shown in Fig. 4B). The left temporal time course is shown in Fig. 4A), with a small non-significant increase in the interaction effect (green line), which peaks after the early window. This is followed by a significant (P_FWE_ = 0.003) second increase that is in the middle window. Clusters are also observed in the frontal area. Fig. 4C) shows the time course for the left frontal cluster. The interaction is significant in the middle window (P_FWE_ = 0.009), with a positive interaction before a reversal of the effect (green line) in the late window, which does not reach significance. The details of the statistical results from SPM are presented in Table 3. This interaction in the middle window suggests that a fronto-temporal network is responsible for linking the local and global effects, and that these two effects (local and global) are not (strictly speaking) independent.

**Table 3.** Statistics for each cluster for the middle windows. Each cluster, named in the first column, is characterized by its peak location in MNI coordinates as shown in the second column, the F-value at the peak (third column), the p-value (fourth column) and the cluster size (last column). The p-value highlights the significant cluster after family-wise error correction, set to an alpha of 0.05.

| Clusters | Peak location<br>(x,y,z) | F[1,119] | P<sub>(FWE)</sub> | K cluster size |
| --- | --- | --- | --- | --- |
| Middle window |  |  |  |  |
| Left frontal | (-46, 20, -10) | 13.11 | $3 \times 10^{-3}$ | 244 |
| Right frontal | (44, 22, -12) | 13.02 | $3 \times 10^{-3}$ | 234 |
| Left temporal | (-46, 20, -10) | 12.45 | $9 \times 10^{-3}$ | 186 |

**Figure 4.** Local-by-global interaction: A) source time-series at the temporal cluster, which is significant in the middle window; B) glass brain of significant clusters for the middle window; C) source time-series at the frontal cluster, which is significant in the middle window. Zero is the onset of the (critical) fifth tone.

### Three way interaction

Finally, the time-series for all conditions and the three-way interaction (local-by-global-by-sedation) with its standard error are shown in Fig 5A). The three way interaction is significant in the late window, with its corresponding significant clusters in the frontal lobe shown in Fig 5D). To characterise the causes of the three way interaction, we explored the two simple effects interactions that constitute it. Specifically, the local-by-global interaction is presented separately for sedation and recovery in Fig 5B) and Fig 5C) respectively. Notably, the local-by-global interaction was significant in the late window when participants had recovered, but not when they were sedated. Indeed, the local-by-global effect (green line) had opposite polarities when sedated and recovered for much of the late window. This difference between sedated and recovered seems to be carried by two properties. Firstly, the LDGD condition terminates more sharply when recovered, and secondly, the LSGD condition has a dramatically higher amplitude when recovered. The former is exactly consistent with the deceleration of the accelerated prediction error reported in [Shirazi2018], suggesting that inferior frontal regions are the source of this shifting neural responsiveness.

**Figure 5.** Three way interaction: A) source time-series for the frontal left cluster of the three - way interaction and the eight conditions involved. B) Local-by-global interaction source time-series for the sedation conditions in frontal cluster. C) Local-by-global interaction source time-series for the recovery conditions in frontal cluster. D) 3D glass brain of the significant clusters for the three way interaction in the late window. E) 3D glass brain of the significant cluster for the local-by-global interaction when recovered.

The latter of these properties (amplitude increase for LSGD) is particularly striking, and important, since the LSGD condition is – in a sense – the most cognitively demanding condition. In particular, there is no bottom-up deviance (as there is in the LDGD condition) signalling an infringement of global regularity. Thus, higher levels in the processing hierarchy effectively need to detect global deviance by the absence of a driving bottom-up prediction error. Our findings suggest that this capacity is realised by inferior frontal regions, consistent with the often discussed role of prefrontal regions in working memory maintenance and update [Polich2007]. Fig 5E) shows the significant clusters for the local-by-global interaction (recovery) in the late window. Statistical results in the late window for the three way interaction and for the Recovery local-by-global simple effect are presented in Table 4, all p-values are highly significant.

**Table 4.** *Statistics of three way interaction for both frontal clusters in the late window, and local-by-global interaction for recovery. Each cluster, named in the first column, is characterized by its peak location in MNI coordinates, as shown in the second column, the F-value of the peak (third column), the p-value (fourth columns) and the cluster size (last column).* The p-value highlights the significant cluster after family-wise error correction, set to an alpha of 0.05.

| Cluster | Peak location<br>(x,y,z) | F[1,119] | P<sub>(FWE)</sub> | K cluster size |
| --- | --- | --- | --- | --- |
| 3 way interaction : late window |  |  |  |  |
| Left frontal | (-46, 20, -10) | 12.21 | $5 \times 10^{-3}$ | 267 |
| Right frontal | (46, 22, -10) | 12.22 | $5 \times 10^{-3}$ | 261 |
| LxG int. for recovery: late window |  |  |  |  |
| Left frontal | (-46, -20, -10) | 17.09 | $< 10^{-3}$ | 516 |
| Right frontal | (44, 22, -12) | 17.24 | $< 10^{-3}$ | 519 |

Additionally, we found a significant effect of sedation in the temporal region, and a significant sedation-by-local interaction in the frontal region, which are presented in the supplementary material (section “Further Sedation Effects”). The sedation-by-global interaction was not found to be significant.

## Discussion

We have presented a source localisation of the neuronal responses evoked by the local-global paradigm – and the effect of sedation with propofol on these responses. In this way, we have addressed how two key cognitive neuroscience theories – predictive coding and the global workspace – are related, and the neurophysiological correlates of this interrelationship.

Importantly, the two theories (predictive coding and Global Neuronal Workspace (GNW)) do already include some related concepts. In particular, prediction has been discussed within the GNW context, for example, the original formulation of GNW [Deheane1998] did discuss “anticipation” and pre-representation. Additionally, and perhaps most notably, [Wacongne2012] presents an important neural instantiation of predictive processing, which simulates the mismatch-negativity. However, we would argue that predictive coding goes further, by providing a full multi-level architecture of brain processing that is theoretically grounded in the mathematics of (generative) Bayesian inference [Friston2010], with concepts such as confidence in (i.e. the precision of) the prediction error to the fore. In this regard, predictive coding provides a full instantiation of the core hypothesis that the cortical architecture is configured to minimize the difference between bottom-up representations, driven from sensory input, and top-down representations of expectations, with this interpretation obtaining at all hierarchical levels. In this sense, there is a clear need to reconcile predictive coding and the Global Neuronal Workspace.

In respect of neural correlates, it is important to acknowledge the constraints associated with our source localisation analysis. In particular, we explicitly placed masks (which were justified by prior precedent; c.f. supplementary material section ‘Subspace selection and mask placement’) to constrain the source reconstruction. In this respect, it is not surprising that we found sources in these a priori regions. However, exactly where those sources fell, especially in the large regions of the frontal and parietal masks, is of interest. Of greater note, is how the MSP algorithm un-mixed variability amongst the regions and how that un-mixing progresses through the time-course of the evoked response. In this respect, a sanity check of our findings is that, as one would expect, early effects are temporal, with a following propagation out from this sensory region to frontal and parietal regions.

The local and global main effects we observed also have considerable face validity. In particular, as shown in Fig 2A, the local effect is detected exclusively in temporal sources in the early window, consistent with sources reflecting the mismatch negativity in auditory cortices [Naatanen2007]. The local effect then engages an (inferior) frontal – (superior) temporal network in the middle window, see figure Fig 2B, which corresponds to the source of the positive rebound to the MMN [Naatanen2007]. This rebound is often related to the P3a, which is known to be generated by frontal sources [Polich2007].

The classic global effect pattern is apparent in the middle and late windows, with what could be considered a prototypical global workspace pattern involving (inferior) frontal – (superior) temporal and parietal regions. Importantly, parietal sources are only present in this contrast, suggesting a unique role for parietal regions in the global workspace. Additionally, there is a clear progression during the global effect from middle to late windows, in which the temporal source wanes, while frontal and parietal sources wax. This suggests a trajectory over time of the global workspace from sensory to encompass association regions. We can highlight four key findings of our analysis:

1. three phases of processing (early, transitional and late);
2. fronto-temporal interaction between local and global in the transitional phase;
3. a failure to detect a main effect of sedation in parietal sources; and
4. a three way interaction between propofol and local and global effects

We elaborate on these in turn.

### 1) Three phases

Fig 6 depicts the neurophysiological realisations of the putative three phases. As discussed previously, the local effect manifests in the early window (early phase) in source space, very much as one would expect – expressed predominantly in superior temporal regions, which include auditory cortices. Additionally, a stereotypical global workspace is present in the late window (late phase). Importantly, our middle window (transitional phase) appears to exhibit qualitatively distinct effects, in terms of the set of sources involved and condition-specific effects exhibited. In particular, the local and global effects only interact in the middle window, suggesting a modulatory exchange between temporal and inferior frontal regions, where the local-by-global interaction was expressed. With a predictive coding and global workspace perspective in mind, one could argue that the three phases can be distinguished according to the following characteristics.

**Figure 6.**
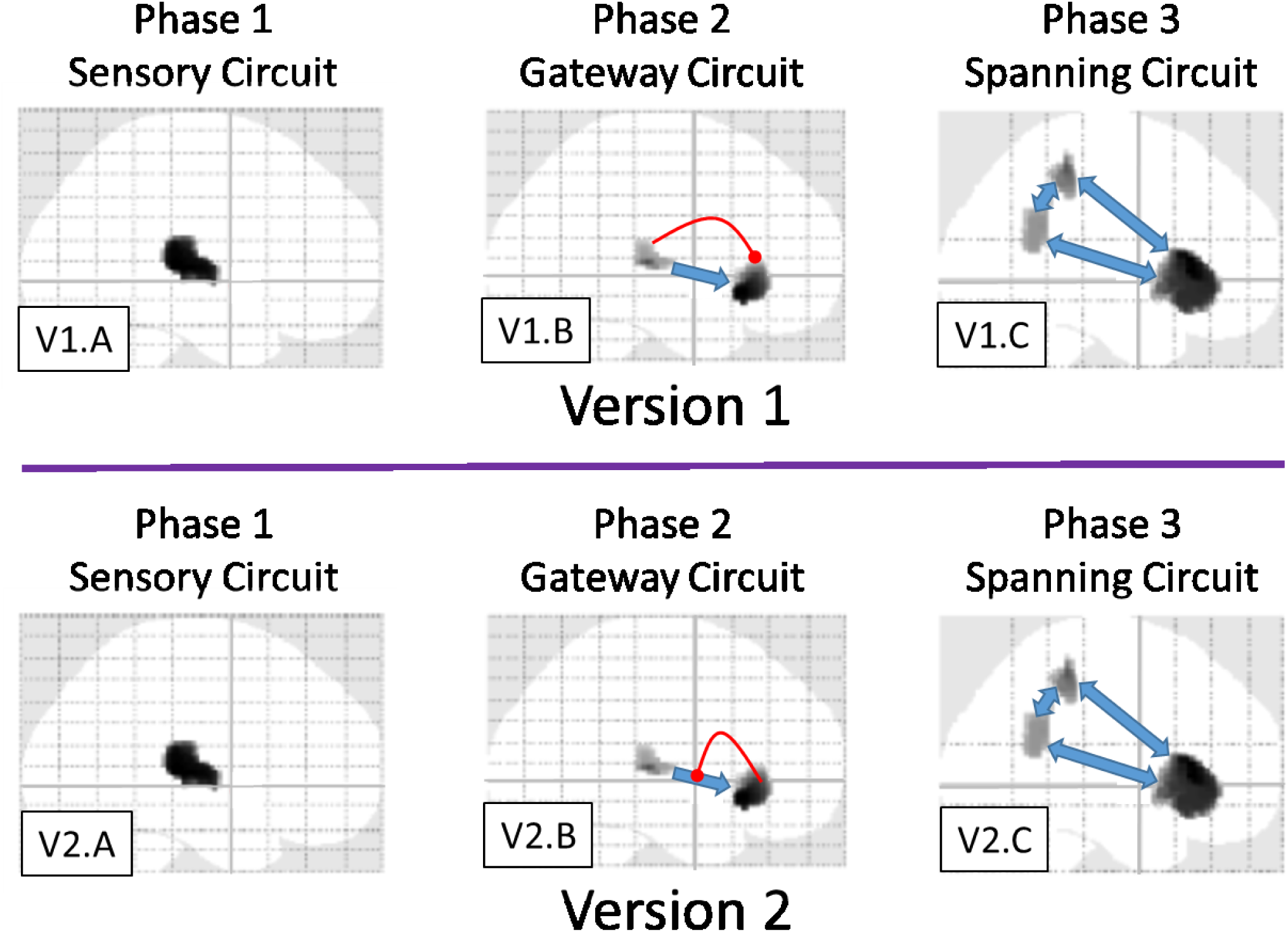
Depiction of three phase theory of local-global processing. Two different versions are presented, which are distinguished by the direction of modulatory activity (c.f. red links). In both versions, phase 1 (c.f. V1.A and V2.A) is restricted to sensory areas, reflecting a sensory prediction error; phase 2 (c.f. V1.B and V2.B) involves interaction between sensory and an inferior frontal region; and phase 3 (c.f. V1.C and V2.C) involves a deep, brain-scale hierarchy, analogous to activation of a global workspace. Importantly, we hypothesise that phase 2 is the transition between phase 1 and phase 3, effectively regulating “ignition” of the global workspace, according perhaps to the precision or priority afforded the ascending (prediction error) signal. The difference between the two versions presented here, with direction of phase two modulation being key, is elaborated in the discussion.

Phase 1 (sensory): activation is restricted to (auditory sensory and perceptual) superior-temporal regions. This phase is expressed over a short time frame, with key responses being consistent with the forward propagation of a bottom-up (sensory) prediction error.

Phase 2 (gateway/transitional): this phase can be hypothesized to engage a mesoscale network (similar to that considered in Phillips et al, 2015, 2016 for the Mismatch Negativity), involving an exchange between (auditory) superior-temporal and inferior-frontal sources. The network features modulatory dynamics, which may implement a priority-based or precision enhancement – consistent with the attentional selection of precise prediction errors. This gateway circuit exhibits a more sustained phasic response than phase 1, but is not meta-stable in the sense of phase 3.

Phase 3 (spanning): this circuit is argued to be macro/brain-scale, spanning the cortical hierarchy. It would naturally be related to the global workspace [Dehaene2011], and would exhibit a prolonged, meta-stable response, as discussed in [King2014].

### 2) Local times global Interaction and the transition phase

to elaborate further, the interaction between local and global in the middle window suggests a multiplicative or modulatory exchange between superior-temporal regions and inferior frontal areas. The former of these elaborates sensory prediction errors that underlie the local effect, and the latter is implicated in higher order processing that integrates over a longer temporal scale.

This network may effectively be a sub-circuit of the brain spanning global workspace, mediating a transitional state; indeed perhaps a proto-workspace. Although further work is certainly required to confirm the hypothesis, it could be argued that the local-by-global interaction in the middle window is suggestive of bidirectional exchanges between levels in a predictive hierarchy, with the multiplicative interaction between levels suggestive of modulation of gain control, which, under predictive coding, could be generated by predictions of precision [Kanai2015].

The Monkey ECoG findings of Chao et al, 2018 sit well with this interpretation. Most importantly, Chao et al, 2018 also highlight a meso-scale circuit between superior temporal areas and prefrontal areas, associated with hierarchical predictive processing. Given the spatial resolution of the source localisation we have performed, there is good alignment between the localisations, with 1) our superior temporal source encompassing both the auditory cortex and anterior temporal sources of Chao et al; and 2) the prefrontal sources of the two studies intersecting (note, monkey pre-frontal cortex is anterior and considerably smaller than human pre-frontal cortex). Additionally, the Dynamic Causal Modelling work on the omission of an expected stimulus (Chennu et al, 2016a) also points to a temporo-frontal circuit with strong feedback influences.

### 3) No parietal sedation effects

given the statistical power available, we did not detect any effects of sedation on parietal areas – while strong effects were found fronto-temporally. This stands against an existing finding of propofol-induced modulation of parietal networks [Schrouff,2011; Uhrig et al, 2016], which we discuss further shortly. From a global workspace perspective, this might seem surprising, since the parietal cortex has been argued to be implicated in conscious experience. This said, with classical statistics, null effects are always difficult to interpret^**3**^, and there remains the possibility that a more highly powered experiment would find an effect at parietal lobe.

Inference though can be less equivocal with regard to the significant effects involving the sedation factor. In particular, we observed a temporal source for the main effect of sedation, see Figs 7A), B) of the supplementary material, which, in the early window, is consistent with the known enhanced N1 in anaesthesia [Ypparila,2004], but for us was also seen in the middle window. Additionally, temporal and frontal sources evidenced a sedation x local interaction; see Fig 7E), F) of supplementary material. This provides some evidence that sedation modulates responses early in the processing pathway. The most interesting effect though was the local by global by sedation effect, which was significant in the late window.

### 4) Three-way Interaction

this was observed at an inferior frontal source; see Fig 5. When we decompose this three-way interaction into its component two-way simple effects – local-by-global when recovered and when sedated – the cause of the three-way is clear. There are two particular aspects to emphasize.

Firstly, the acceleration of the global deviant response by the coincidence of local deviance (the double surprise acceleration effect [Shirazi2018]), is evident at the inferior frontal source both when sedated and when recovered, with this acceleration being stronger when recovered. This is apparent in the sharper onset and offset of the LDGD condition when recovered compared to sedated; see Fig 5B) and C). This suggests that the deceleration of the accelerated prediction error described in [Shirazi2018], can be localised to inferior frontal regions.

Secondly, the most striking feature driving the three way interaction at frontal in the late window is the dramatically higher LSGD condition when recovered than when sedated; see Figs 5B), C). Importantly, the LSGD condition is most dependent upon long-term temporal integration. In particular, this global deviance is not marked by a sensory prediction area (since it arises during a local standard quintuple). Thus, the deviance is not initiated by a strong bottom-up signal (i.e. a local prediction error). That is, it is a pure global deviance condition, with its detection intrinsic to higher hierarchical levels.

It seems then that sedation impairs this capacity to detect deviance intrinsically at higher levels, at least at inferior frontal sources. This finding is in many respects consistent with the intent of the local-global task; i.e., to differentiate processing that requires temporally sustained integration over an extended period of time, and the role of consciousness in this temporally extended evidence accumulation. Thus, our findings provide suggestive evidence that, in respect of the action of propofol, reduced awareness diminishes long duration processing of temporal integration, supported by inferior frontal sources. In terms of predictive coding, this finding is consistent with a reduction in the precision of ascending prediction errors. This follows because precision corresponds to the rate of evidence accumulation (Hesselmann, Sadaghiani et al. 2010, FitzGerald, Schwartenbeck et al. 2015). In other words, belief updating in response to precise prediction errors converges more quickly than in the setting of imprecise prediction errors – or a pharmacological reduction in gain of responsiveness of populations encoding prediction errors. This particular effect of propofol would endorse the hypothesis we highlighted earlier that ignition depends upon a phase transition that itself rests upon attentional selection of ascending prediction errors that is mediated by the modulatory effects of predicted precision.

## Sedation

The “sedation state” that we are exploring involves placing the participant at the fringe of consciousness, which may correspond to a (weakened, but) active bottom-up stimulus strength and lower top-down attention. In this regard, it may be comparable to the “pre-conscious” state in [Dehaene2006]. However, it is notable that in our data, sedation reduces, although seems not to eliminate, the sensory response, e.g. see supplementary material section “Further Sedation Effects”, where Figure 7A,B,C shows a clear sedation effect at sensory areas. Because Event Related Potentials are averages across many trials, it is always difficult to know whether the reduction of a component is due to a consistent reduction at the single-trial level, or increased variability in the response. Thus, it is possible that the reduction in sensory components with sedation that we observe are caused by intermittent activation, in which sensory input is extremely weak, even absent, on some trials, and strong, presumably with the global workspace fully engaged, on others. Resolving these competing explanations awaits further investigations.

Our findings on the effect of propofol sedation resonate with a number of previous findings. Firstly, Boly et al [Boly2012] suggest that patients with reduced levels of consciousness (i.e., vegetative state) exhibited a reduction in effective connectivity for a descending extrinsic (between source) connection from inferior frontal to superior temporal regions. This link was found absent during a mismatch negativity paradigm, very similar to the local component of the local-global task. While we are limited in our capacity to decompose our temporal region anatomically, and thus to directly implicate superior temporal regions beyond primary auditory cortex, we have found effects of sedation in (superior) temporal/inferior-frontal regions. Although, care should certainly be taken in relating two different forms of reduced awareness (vegetative state and sedation).

Uhrig et al, 2016 present an impressive fMRI study of the effects of anaesthesia on brain responses to the local-global effect in monkeys, in which they report a reduction in prefrontal responses to global deviance during sedation. Additionally, Nourski et al, 2018 observed a striking abolition of pre-frontal responses to global deviance with sedation. As previously discussed, we observe similar effects, although our findings are within the context of our local-by-global interaction enabling us to clarify the non-additive relationships between local and global levels. Additionally, Uhrig et al observed an effect of sedation on the local effect, which is consistent with the local x sedation interaction we report in the Supplementary Material (subsection Further Sedation Effects).

Where there is some inconsistency with our findings is in respect of parietal sources. Uhrig et al, 2016 identified reduced global deviant responses parietally with anaesthesia, however, we failed to find any significant interaction effects involving sedation and global at parietal sources. A failure to reject the null does not, of course, enable its affirmation, leaving this question not definitively answered. Although, it should be noted that, (in a Psycho-Physiological Interaction) for the global effect, Uhrig et al did observe a residual context-dependent coupling between auditory areas and intra-parietal sulcus during moderate Propofol sedation, suggesting that, even in their study, sedation did not fully obliterate parietal responses for the global effect.

In summary, although there remains considerable uncertainty, the key ERP components we observe are (increased) responses to unexpected/deviant stimuli, which would naturally be considered a signalling of prediction error. In this context, we can give a more specific candidate interpretation of the effects of sedation: predictive coding suggests that the feed-forward error signal reflects a prediction error weighted by its precision or confidence afforded that error signal. Confidence in our context would be driven by an assessment of the level of irreducible sensory prediction error – or by descending predictions of precision based upon belief updating higher in the hierarchy (see Figure 6). The two key effects of sedation that we observe are (1) a reduction in amplitude and (2) a slowing of the evoked response to deviant stimuli, which are particularly pronounced with global deviance. Indeed, this is marked in the evoked response to the simultaneous confounding of expectations at multiple hierarchical levels; namely local and global. A likely candidate for both of these reductions – in amplitude and in speed – would be reduction of gain/precision [Kanai2015], which would induce a broad loss of responsiveness.

## Top-down or bottom-up ignition?

Our source localisation implicates a superior-temporal – inferior-frontal circuit in this modulation of response by sedation. Although – on the basis of the findings presented here – we cannot be certain of the direction of modulatory influence (descending or ascending) in this circuit. This is reflected in the two versions of the three phase theory presented in Fig 6.

[Shirazi2018] proposed that the acceleration of the global response – due to coincidental local deviance – is caused by a feed-forward modulation from the local effect circuit onto the global effect circuit. This is the direction of modulatory influence presented in Fig 6, version 1, see panel V.1B. This might be considered an explanation of the acceleration by double surprise that sits most easily with the simplest line of temporal causation, since registration of local deviance would naturally be considered to precede registration of global deviance.

However, the other direction cannot be excluded. That is, it could be that a weakening of the modulatory influence of inferior-frontal on superior-temporal areas is what drives the slowed and attenuated responses observed when sedated. Such an explanation would be consistent with the second version of our three phase theory presented in Fig 6, where modulation is mediated in a feedback direction (i.e., descending predictions of precision), see panel V2.B. From a predictive coding perspective, as previously suggested, a potential explanation of the effect of sedation is that it reduces the precision of sensory prediction errors, potentially carried by a feedback link from inferior frontal to temporal regions, as per Fig 6 (V2.B). This would effectively attenuate the gain on the ascending transmission of prediction errors. As noted in the introduction, this formulation of the transition from local to global processing (i.e., ignition of the global workspace) provides a graceful synthesis of predictive coding and global workspace theory that is grounded in neurophysiology (via gain control and neuromodulation) – while at the same time speaks to psychological concomitants of conscious processing (via attentional selection that accompanies perceptual synthesis over extended periods of time).

## Acknowledgements

We would like to thank Guillaume Flandin for very valuable discussions of SPM analyses.

## Footnotes

1 *The term accelerate is primarily used metaphorically, although, the advanced P3 component that we identified in [Shirazibeheshti2018] (see figure 5 LxG panels in that paper) does in fact exhibit an acceleration in the mathematical sense. That is, the tangent to the curve (the velocity) changes rapidly with time, meaning that the rate of the rate of change with time (i.e. the acceleration) is bigger in the advanced P3 (the LDGD condition).*

2 https://www.sciencedirect.com/topics/medicine-and-dentistry/ramsay-sedation-scale

3 Ideally, one would like to perform a Bayesian analysis to find evidence for the null. However, it is not currently clear how to formulate a Bayes Factor that accurately reflects the inferential steps involved in a family-wise error corrected neuroimaging analysis.

## Notes

https://www.repository.cam.ac.uk/handle/1810/252736

